# *In vitro* direct regeneration and *Agrobacterium tumefaciens* mediated *in planta* transformation of *Ocimum sanctum L.*

**DOI:** 10.1101/2022.01.31.478449

**Authors:** Sana Khan, Zakir Husain, Laiq ur Rahman

**Author notes:** Corresponding Author Dr. Laiq ur Rahman, Senior Principal Scientist, Plant Biotechnology Division, CSIR-Central Institute of Medicinal & Aromatic Plants (CIMAP), Lucknow-226015 (U.P.) INDIA. Academy of Scientific and Innovative Research (AcSIR), CSIR-Central Institute of Medicinal and Aromatic Plants (CIMAP), Lucknow, India.

## Abstract

*Ocimum sanctum* is a multipurpose herb with highly significant medicinal properties. An *in vitro* direct regeneration protocol for propagation of a valuable medicinal plant *Ocimum sanctum*, using petiole explants has been successfully developed. The protocol employed regeneration of shoots directly, without any intervening callus using Murashige and Skoog (MS) medium fortified with 3 mg L^-1^ BAP + 1 mg L^-1^ NAA. The maximum regeneration frequency of 98% with 9.6 shoots per explants was achieved. *Agrobacterium tumefaciens* mediated genetic transformation (ATMT) protocol (transient and stable) was established using LBA4404 strain harboring pBI121 with *uid-A* reporter gene and neomycin phosphotransferase (*npt-II*) as selection marker. The putative transformants were screened on MS with 50 mg L^-1^ kanamycin and subsequently rooted on the half-strength MS medium. The confirmation was done *via* polymerase chain reaction (PCR) using *npt-II* and *gus-A* gene-specific primers. The maximum stable transformation frequency 70% ± 0.35. Thus, it is apparent that the established *in vitro* direct regeneration and ATMT method was suitable for integrating novel genes and modulating the metabolic flux for obtaining desired agronomic trait *in planta*.

## Introduction

*Ocimum sanctum L*. (*O. tenuiflorum*) is a valuable sacred medicinal, and aromatic plant commonly called as “holy basil” aka Tulsi.^1^ It has been used in traditional, ayurvedic, Unani system of medicinal systems. ^2,3^ The sacred tulsi herb is known as ‘The Incomparable One’, ‘The Queen of Herbs’ for more than a decade and worshipped for over 3000 years.^4^ The bioactive phytoconstituents derived from *O. sanctum* have a rich repository that inherits several potential therapeutic and pharmaceutical utilities. Several phytochemicals such as eugenol (1-hydroxy-2-methoxy-4-allylbenzene), euginal, urosolic acid,^5^ carvacrol limatrol, caryophyllene, and anthocyans^6^ are the principal ingredients with therapeutic and traditional values since ages. The essential oil of *O. sanctum* showed efficacies as cardioprotective, anti-fertility, anti-diabetic, anti-microbial, anti-cancer, anti-fungal, analgesic, anti-spasmodic, and adaptogenic actions, and so on.^6^

Despite having enormous pharmacological utilities, the underpinning pathways in *O. sanctum* remains insufficiently understood due to the lack of genetic transformation protocol. As a prelude to increasing the production of desired secondary metabolites through targeted breeding programs, the transcriptomic and genomics data needs to be explored and thus an efficient regeneration and transformation protocol is urgently required. Yet, few of the transformation protocols have been reported such as (i) in *O. gratissimum* where only 20% ± 0.7 stable transformation frequency was achieved^7^; while (ii) in *Ocimum basilicum* and *O. citrodonum*; only transient expression system was established by Deschamps and Simon.^8^

Therefore, for the first time, an efficient and rapid direct *in vitro* regeneration and ATMT protocol (transient as well as stable) has been reported in ‘*O. sanctum’(cim-ayu)*. The established protocol is highly efficient and reproducible in the rapid regeneration of transgenic plantlets with a stable expression system. The method could be used as a potential tool in developing improved varieties with high therapeutic and economic value. Isolation and characterization of novel genes would also be done in order to modulate the desired transcriptional/translational regulons, which paves the way for further advancement in engineering of agronomic traits in related species.

## Materials and Methods

### Establishment of *in-vitro* culture and Direct Regeneration in *O. sanctum*

Seeds of *O. sanctum* were collected from the National gene bank of medicinal and aromatic plants (NGBMAP) at CSIR-CIMAP, Lucknow. The seeds were rinsed with 70% ethanol followed by surface sterilization using 0.1% of HgCl_2_ for 60s. Treated seeds were further rinsed thoroughly with double distilled autoclaved water and placed on basal MS medium^9^ supplemented with 3% (w/v) sucrose and 0.8% (w/v) agar. The pH of the medium was adjusted to 5.8 ± 0.5 prior to autoclave at 121 psi for 15 min. The seedlings were allowed to grow into full plantlets.

After 4–5 week of establishing *in vitro* culture of *O. sanctum*, several explants such as hypocotyls, cotyledonary leaf and petiole were excised and used for direct *in vitro* regeneration studies. The explants was placed on modified MS medium supplemented with BAP (0.5 mg/L to 3.0 mg/L), in combination with NAA 0.5 mg/L to 1.5 mg/L. The data was recorded after 4–5 weeks of inoculation of culture on the most responsive medium termed as shoot multiplication medium (SMM). All cultures were maintained under cool fluorescent light (3000 lux) with 16 L: 8 D-h photoperiods at 25 ± 5ºC. Afterward, *in vitro* proliferated shoots (approx. 1.5 cm) were excised and shifted to hormone-free MS medium for elongation and ½ MS for root induction, respectively. The rooted plantlets having 5–6 leaves were thoroughly washed to remove agar and acclimatized in Hoagland’s medium for one week and hardened into small pots [vermiculite and sand (1:1)]. The plantlets were shifted to glass house for further growth and development. Out of several explants, petioles were used for further genetic transformation studies as the best regeneration response was achieved from petioles.

### Optimization of Kanamycin concentration for selection of putative transformants

To investigate the optimal concentration of kanamycin for the screening of putative transformants, a sensitivity test for shoot regeneration was studied in *O. sanctum* using petioles. The explants were inoculated on modified MS medium supplemented with 3.0 mg/L BAP, 1.0 mg/L NAA (best suited SMM), and different kanamycin concentrations (10–60 mg/L). The petioles that were cultured on SMM without kanamycin served as a control (0 mg/L). The selection of the putative transformed regenerated shoot and data was recorded after 4-5 weeks of culture.

### Optimizing parameters for *Agrobacterium tumefaciens* mediated genetic transformation protocol in *O. sanctum*

#### *Agrobacterium tumefaciens* strain and transformation vector

*A. tumefaciens* strain LBA4404 harboring binary vector pBI121 with *uidA* gene (β-glucuronidase) as marker and neomycin phosphotransferase-II (*nptII) as* selection marker gene was used for genetic transformation study in *O. sanctum*. The *gus A* and *npt II* gene present on T-DNA were driven under the control of cauliflower mosaic virus (CAMV) 35S promoter and Pnos promoter (nopaline synthase promoter) respectively.

#### *Agrobacterium tumefaciens* mediated genetic transformation

*A. tumefaciens* was grown in YEP minimal medium containing 25 mg/L rifampicin, 50 mg/L kanamycin, and 250 mg/L streptomycin at 28ºC for overnight, at 200 rpm on a rotary shaker. *A. tumefaciens* cells were harvested [OD(600) 0.6] by centrifugation at 5,000 rpm for 10 min, and pellet was resuspended in equal volume of MS liquid medium. The explants were immersed in an agro bacterial infection medium in the presence of AS (0-400 µM), Tween-20 (0.001-1%) and kept on a shaker (100rpm) for 0-60 min. The infected explants were then blotted dry onto a sterile filter paper and co-cultivated on CCM for 48 hrs in the dark. Afterward, the explants were transferred onto the SMM fortified with 3 mg/L BAP, 1 mg/L NAA, 50 mg/L kanamycin and 250 mg/L cefotaxime for direct regeneration at 25 ± 5 ºC.

### Histochemical GUS assay

The putative transformed, regenerated shoots were assessed for transient as well as stable gus expression. The explants were excised from the transformed plant using sterile blade. For transient gus expression, the treated explants were checked just after the infection, while stable gus expression was analysed after 3 months of culture with kanamycin-resistant regenerated transgenic shoots. The explants were incubated at 37ºC overnight (in the dark) in 1mM 5-Bromo-4-chloro-3-indoyl β-D-glucuronide-cyclohexylammonium salt (X-Gluc), 0.1% Triton X-100, 0.1mM potassium ferricyanide, 0.1mM potassium ferrocyanide, 0.1 M sodium phosphate buffer (pH 7.0). Following incubation, the chlorophyll of the explants was removed using 70% (v/v) ethanol.

### Molecular characterization

Genomic DNA was extracted from young leaves of transformed and non-transformed (control) plants of *O. sanctum* using CTAB protocol.^10^ Polymerase chain reaction (PCR) amplification was carried out with gene-specific primer for both *nptII* and *gus-A* gene [F: 5’ATCAGGATGATCTGGACGAAGAG3’ and *npt-II* R:5’CAAGCTCTTCAG CAATATCACG3’ and F: 5’ACTGTAACCACGCGTCTGTTGAC3’ and *gus-A* R: 5’ TGTTTGCCTCCCTGCTGCGG 3’]. Amplification for *npt-II* was carried out at 94ºC for 5 min as denaturation, 32 subsequent cycles of 94ºC denaturation for 30s, 56ºC for 30s as annealing, 72ºC for 30s, and elongated at 72ºC for 5 min as final extension. Likewise, for *gus-A*, 94ºC for 5 min as denaturation, 35 subsequent cycles of 94ºC denaturation for 1 min, 52ºC for 30s as annealing, 72ºC for 45s; and elongated at 72ºC for 10 min. Plasmid of pBI 121 was used as a positive control. The amplified products were resolved by electrophoresis on 1.2% (w/v) agarose gel.

### Statistical Analysis

Completely randomized experimental design was followed during the experiment. Each treatment experiment consisted of 30 explants per petriplate with three replicates per treatment. Data of shoot regeneration were analyzed using one-way analysis of variance (ANOVA). The significance of difference was observed (P < 0.05).

## Results and discussion

### Direct Regeneration and Multiple Shoot Induction in *O. sanctum*

To develop a suitable expression system, it is a prerequisite to have efficient regeneration and genetic transformation methods that must be repeatable and easily reproducible. The success of *Agrobacterium tumefaciens* mediated genetic transformation and the expression frequency of any gene is highly depends upon regeneration frequency, which provides a pavement to recover multiple micro shoots as well as transgenics. Therefore, prior to an experiment, it is highly recommended to have a standardized protocol to achieve appropriate results.

In order to induce direct regeneration in *O. sanctum*, several explants (leaf, internode, petiole, hypocotyl etc.) were placed on full strength basal media fortified with a number of combinations of PGRs (Fig. 1A-O). However, the direct regeneration and multiple shoot induction/proliferation were successfully achieved maximally through petiole, followed by hypocotyl, and leaf (Fig. 2A-C). In the present study, *de novo* axillary/apical shoot regeneration was achieved successfully from petiole are in concordance with those for Populus, Jatropha, and pigeon pea^11-14^. *In vitro* petioles were incubated with MS having different concentrations of BAP (cytokinin) and NAA (auxin). The concentration of 3 mg/L BAP and 1 mg/L NAA was found to be most effective among all combinations (Fig. 2D,E). The same combination yielded the maximum number (9.6) of regenerated shoots per explant with the highest shoot regeneration rate i.e., 98%. At the same time, further change (either increase or decrease) declines both regeneration rate and quantity of buds originating from single explants (Table 1). The obtained result indicates that the efficiency of *de novo* shoot induction was influenced significantly at different concentrations. An exogenous supply of cytokinins (with or without auxin) stimulates axillary shoot formation. ^15^

**Fig. 1.**
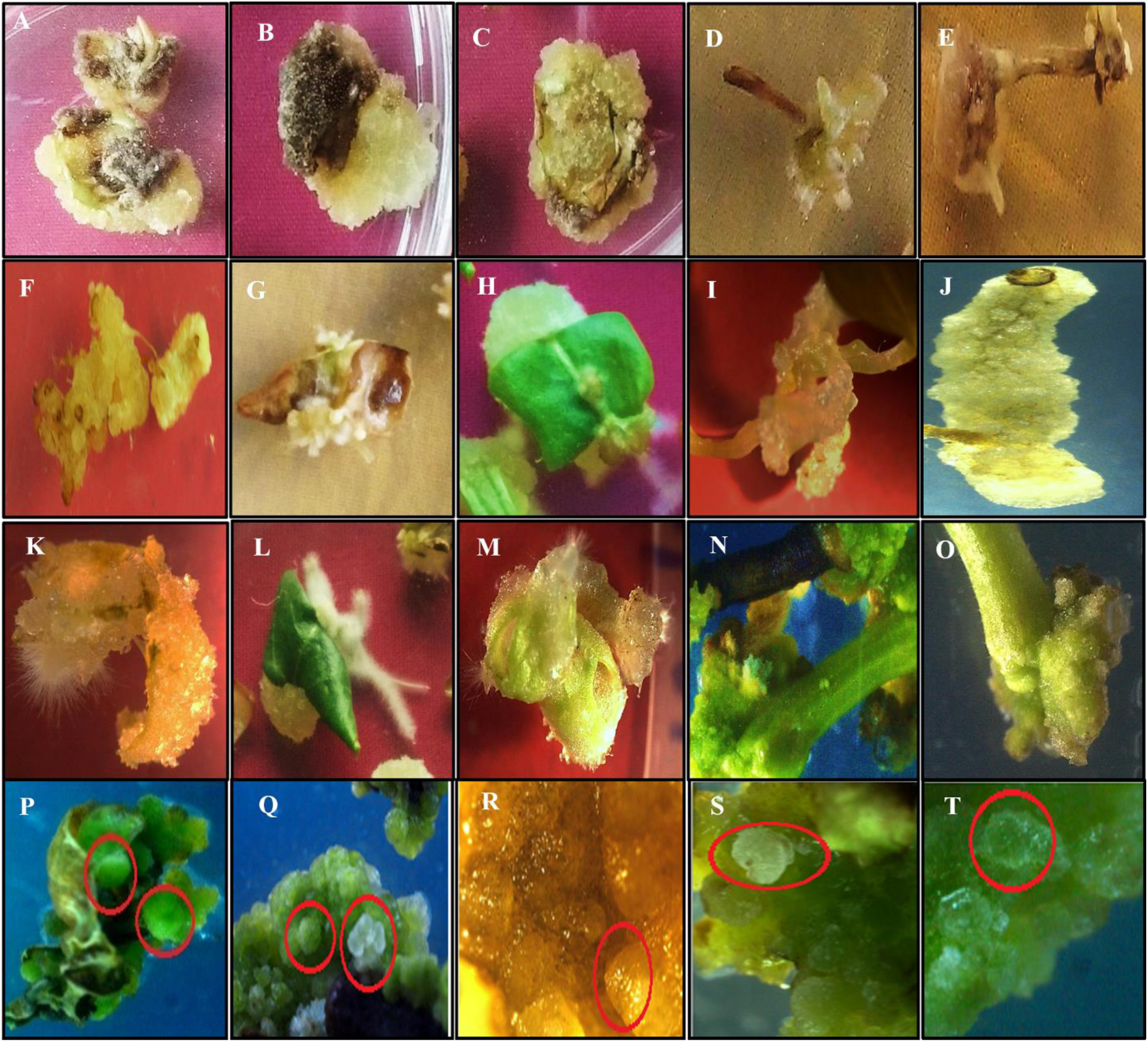
Influence of several different combinations of plant growth regulators on various explants for direct regeneration and multiple shoot induction on *O. sanctum*.

**Fig. 2.**
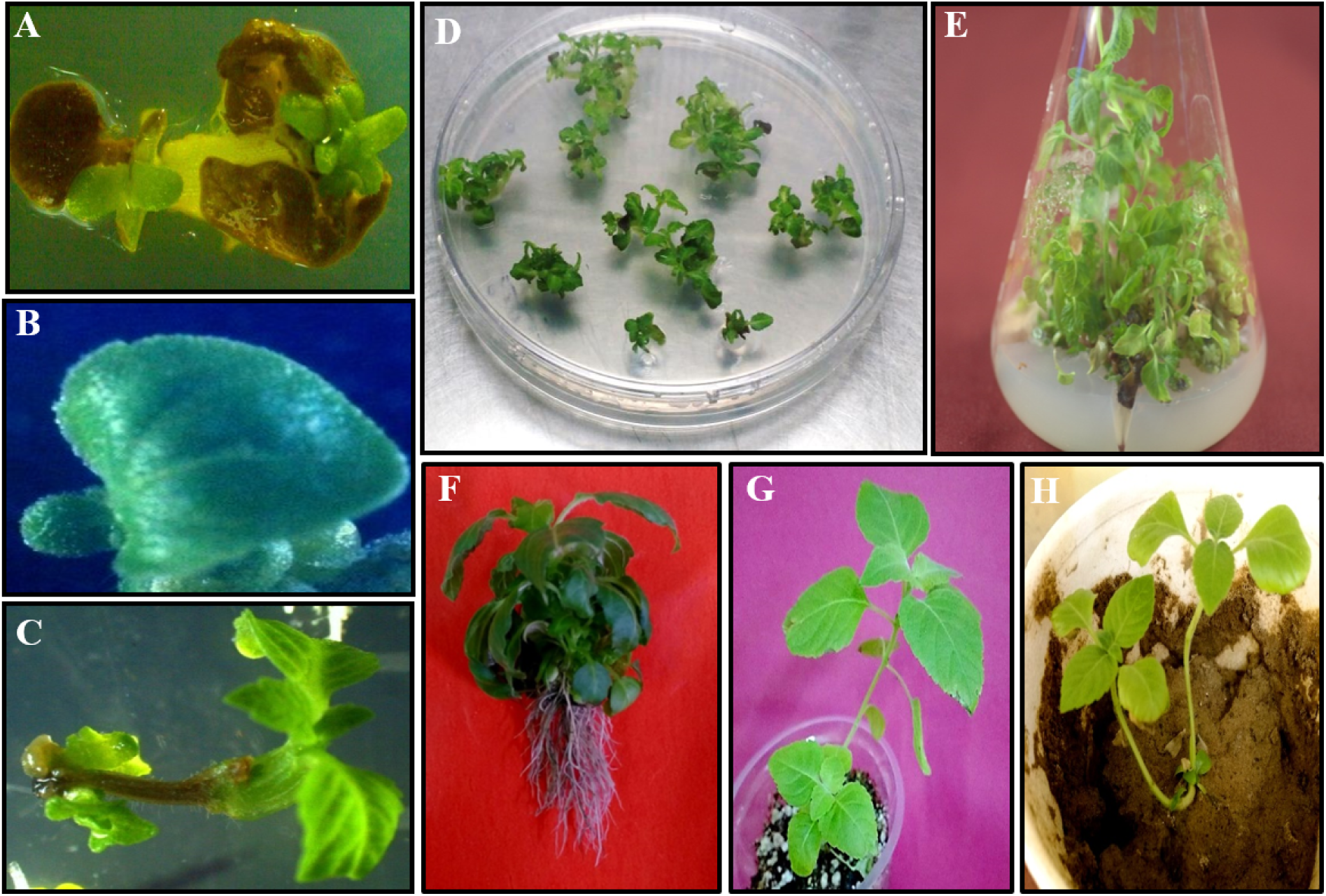
Direct regeneration, Multiple shoot proliferation, and Root induction in *O. sanctum*. **(A)** Shoot regeneration from hypocotyl, **(B)** Shoot regeneration from cotyledonary leaf, **(C)** Shoot regeneration from petiole, (**D**) Petioles showing direct regeneration and multiple shoot induction, (**E**) *In-vitro* culture showing multiple shoots, (**F**) *In vitro* regenerated plantlets with healthy root system (**G**,**H**) Hardening and Acclimatization.

**Table 1.** Effect of BAP and NAA concentrations on direct shoot regeneration from different explants of *O. sanctum*

| MEDIA<br>COMPOSITION<br>(MS+BAP+NAA)<br>mg/L | Explant |  |  |  |  |  |
| --- | --- | --- | --- | --- | --- | --- |
|  | Hypocotyls |  | Cotyledonary Leaf |  | Petioles |  |
|  | No. of | Response | No. of | Response | No. of | Response |
|  | shoots/explant | (%) | shoots/explant | (%) | shoots/explant | (%) |
| 0.5+0.5 | 1 ± 0.7 | 10.96 ± 0.85 | 1.033 ± 0.95 | 7.36±0.27 | - | - |
| 0.8+0.5 | $1.61 \pm 0.81$ | $9.16 \pm 0.87$ | $1.47 \pm 0.49$ | $6.70 \pm 0.29$ | - | - |
| 1+0.5 | $1.13 \pm 0.32$ | $9.6 \pm 0.92$ | $2.0 \pm 0$ | $10.7 \pm 0.65$ | $3.66 \pm 0.28$ | $30 \pm 1.00$ |
| 1.5+0.5 | $0.5 \pm 0.3$ | $13.63 \pm 1.01$ | $1.28 \pm 0.62$ | $6.3 \pm 0.33$ | $5.0 \pm 1.00$ | $46.6 \pm 0.57$ |
| 2+0.5 | $0.43 \pm 0.15$ | $15.8 \pm 0.50$ | $1.68 \pm 0.60$ | $4.46 \pm 0.43$ | $7.6 \pm 1.52$ | $73.3 \pm 2.08$ |
| 2.5+0.5 | $0.86 \pm 0.41$ | $9.43 \pm 0.91$ | $2.17 \pm 0.18$ | $7.16 \pm 0.41$ | $7.0 \pm 1.73$ | $55 \pm 1.15$ |
| 3+0.5 | $1.06 \pm 0.25$ | $6.33 \pm 1.09$ | $1.13 \pm 0.28$ | $1.8 \pm 0.31$ | $8.33 \pm 0.57$ | $63.3 \pm 1.15$ |
| 1+1 | $1.18 \pm 0.16$ | $9.48 \pm 0.46$ | $2.08 \pm 0.86$ | $4.28 \pm 0.02$ | $2.60 \pm 1.52$ | $15 \pm 0.50$ |
| 1.5+1 | $1.32 \pm 0.58$ | $8 \pm 2.06$ | $0.91 \pm 0.13$ | $2.20 \pm 0.29$ | $1.60 \pm 0.28$ | $18 \pm 0.51$ |
| 2+1 | $2.46 \pm 0.80$ | $3.33 \pm 0.327$ | $2.82 \pm 0.28$ | $1.87 \pm 0.22$ | $2.70 \pm 1.15$ | $20 \pm 0.76$ |
| 2.5+1 | $2.16 \pm 0.22$ | $3.73 \pm 0.380$ | $2.22 \pm 1.30$ | $4.37 \pm 0.66$ | $7.33 \pm 2.08$ | $70 \pm 1$ |
| 3+1 | $1.58 \pm 0.52$ | $5.76 \pm 0.807$ | $1.66 \pm 0.57$ | $2.63 \pm 0.28$ | $9.67 \pm 0.57$ | $98.3 \pm 0.28$ |
| 1+1.5 | $0.94 \pm 0.37$ | $3.3 \pm 0.38$ | $1.106 \pm 0.83$ | $3.40 \pm 0.30$ | - | - |
| 1.5+1.5 | $0.89 \pm 0.21$ | $3.33 \pm 0.57$ | $1.25 \pm 0.69$ | $7.35 \pm 0.64$ | - | - |
| 2+1.5 | $1.81 \pm 1.27$ | $4.83 \pm 0.50$ | $2.19 \pm 1.32$ | $9.50 \pm 0.89$ | $2.30 \pm 2.08$ | $53.3 \pm 0.28$ |
| 2.5+1.5 | $1.35 \pm 0.90$ | $6.63 \pm 0.57$ | $3.53 \pm 2.49$ | $3.96 \pm 0.26$ | $5.30 \pm 2.09$ | $70 \pm 1.0$ |
| 3+1.5 | $1.64 \pm 0.51$ | $9.56 \pm 0.93$ | $2.34 \pm 1.11$ | $3.06 \pm 0.55$ | $8.4 \pm 1.15$ | $73.33 \pm 1.15$ |

However, when BAP and NAA concentration increased from 1 to 3 mg/L BAP and 0.5 to 1 mg/L NAA, significant variation in regeneration percentage of regenerated axillary shoots was observed. Similarly, the number of induced apical shoots per explant was also affected accordingly. On further increase in the concentration of hormones, it became supra-optimal for the petioles to differentiate and showed hyper-hydricity without any further increase in the number of regenerated shoots. It strongly agrees with the results obtained during the present study that elevating the cytokinin (BAP) in SMM induced de novo apical shoot formation^16,17,18^ but at the same time, reduced cytokinin resulted in less number of regenerated shoot (Table 1).^19^ Additionally, it has been noted that a high concentration of cytokinins inhibits shoot regeneration rate and resulted in poor shoot with symptoms such as rosette shoots and vitrification. ^20,21^

Therefore, the present study highly corroborates to previous findings, in which the existence of crosstalk between auxin and cytokinin signalling showed to interact both synergistically and antagonistically during the development of axillary meristems.^22-26^ Furthermore, the results achieved were in line with previous reports that showed significant achievements in terms of direct regeneration.^7,11,27^ De novo plant regeneration from petiole was achieved successfully through synergistic interaction of hormones (cytokine and auxin) directly.^28,29^ Additionally, formation of embryo like structures were observed but did not germinate into plantlet (Fig. 1P-T). An efficient and highly reproducible method for continuous high-frequency shoot induction and multiplication along with plantlets production in *O. sanctum* was developed. The present study could be helpful in studying the crosstalk between hormonal-induced and endogenous programs that have not well understood yet. Also, the protocol is highly proficient and reproducible to induce large scale clonal and micro propagation in *O. sanctum*.

### Root induction of *in vitro* regenerated shoots

Green regenerated microshoots (1-1.5 cm) were transferred to full and half-strength MS medium, along with MS having varied concentrations of IBA or NAA (0.1 −1.5 mg/L) for root induction. After 4-5 weeks of culture, it was observed that half-strength MS basal medium significantly influenced the response and frequency of root induction than that of MS medium fortified with either IBA and NAA and full strength MS medium as shown in Table 2. Results showed that ½ MS basal medium could effectively stimulate the initiation and growth of roots, inducing the best effect and the highest root induction rate (85 ± 0.35%). IBA and NAA were found to be efficient in the root induction *in planta*.^30^ However, in our study, supplementing with either IBA or NAA had not shown any significant result. The results obtained are in conformity with the previous results^31^ (Table 2). The roots regenerated on 1/2 medium were healthy, while those on full-strength MS medium and IBA /NAA were tender, slim, and eventually turned to black. The root formation was initiated after 2-3 weeks of subculture, and the regenerated plantlets were observed to have well-established root system (Fig. 2F). These plantlets were further transferred and hardened into small pots containing vermiculite and sand (1:1) for acclimatization and proper growth (Fig. 2G,H).

**Table 2.** Influence of NAA or IBA, ½ MS and MS on rooting of *in vtiro* derived shoots of *O. sanctum*

| MEDIA COMPOSITION | No. of roots/explants | Response (%) |
| --- | --- | --- |
| Half strength MS | $7.6 \pm 0.27$ | $85 \pm 0.35$ |
| Full strength MS | $4.4 \pm 0.3$ | $60 \pm 1.5$ |
| MS + 0.1 NAA | $4.0 \pm 0.49$ | $40 \pm 3.44$ |
| MS + 0.5 NAA | 4.46 ± 0.38 | 40 ± 2.08 |
| MS + 1 NAA | 4.1 ± 0.25 | 43 ± 1.20 |
| MS + 1.5 NAA | 3.5 ± 0.17 | 37 ± 0.8 |
| MS + 0.1 IBA | 3.7 ± 0.24 | 45 ± 0.17 |
| MS + 0.5 IBA | 3.2 ± 0.1 | 58 ± 0.15 |
| MS + 1 IBA | 3.28 ± 0.31 | 42 ± 0.23 |
| MS + 1.5 IBA | 2.4 ± 0.29 | 48 ± 0.06 |

#### *Agrobacterium tumefaciens* mediated genetic transformation studies in *O. sanctum*

### Optimization of selection conditions in *O. sanctum*

The sensitivity of the explants to kanamycin was assessed by adding varying concentrations of kanamycin (10-60 mg/L) in SIM and visually checking the regeneration of explants after 3-4 weeks of culture. The petioles cultured on MS medium without kanamycin (0 mg/L) were healthy, and green showed the highest regeneration responses. The use of antibiotic selection marker like kanamycin included within the same vector along with the desired genes provides an efficient and convenient screening system. In the present study, explants at 30 mg/L kanamycin showed a regeneration frequency of 70% ± 0.76 with healthy green shoots. However, on further increase (40 mg/L); the regeneration frequency was declined (13% ± 0.28). Meanwhile, the untransformed regenerated shoots inoculated on 50-60 mg/L kanamycin have shown complete bleaching along with the induction of albino microshoots that eventually turned to brown (Fig. 3B). The result can be correlated with previous findings where kanamycin was used as a successful screening marker.^32,33^ Therefore, for present study, 50 mg/L of kanamycin concentration was optimum as the minimum inhibitory concentration to remove pseudo-escapes and effective screening of putative transformants.

**Fig. 3.**
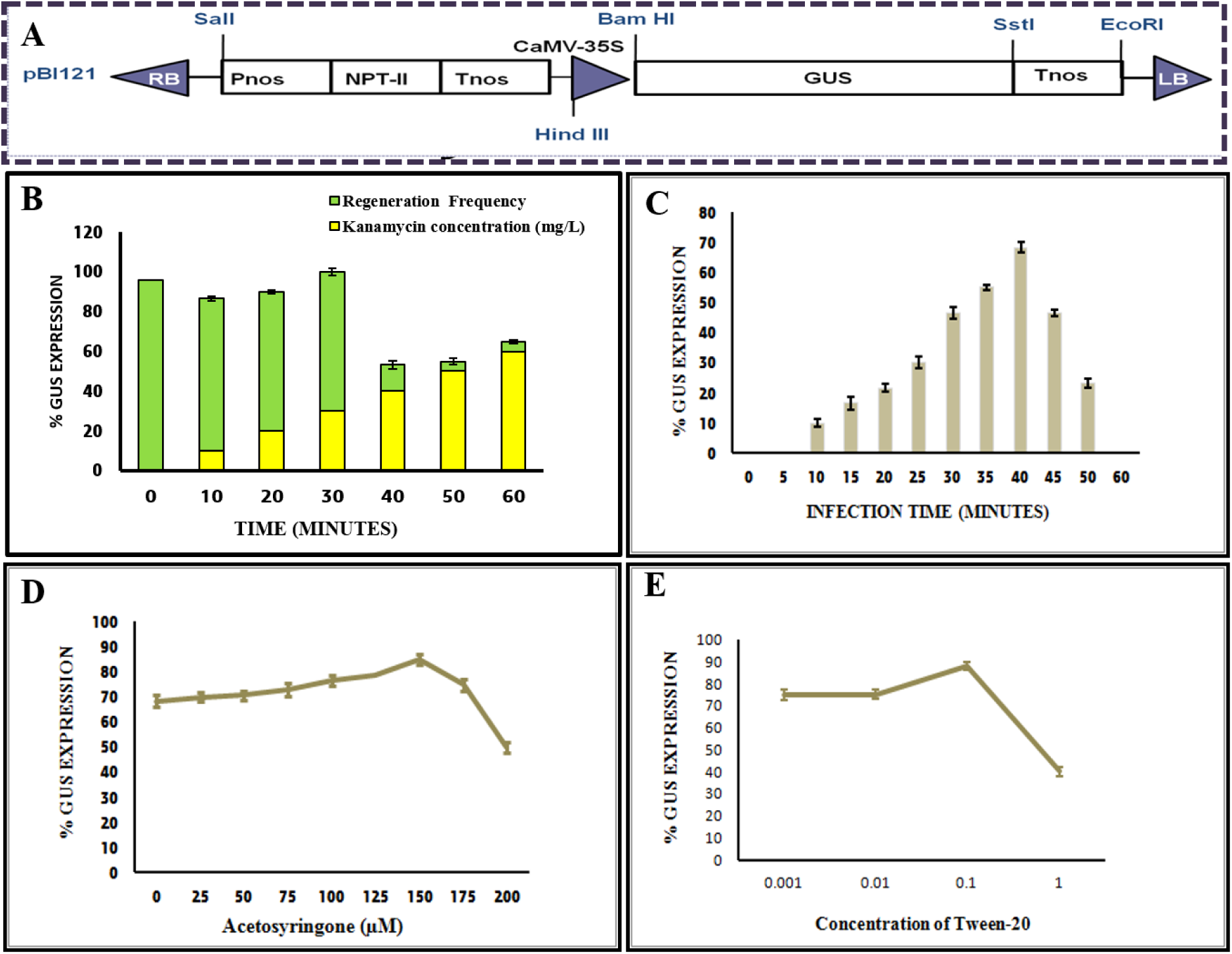
T-DNA construct map and optimized parameter for *A. tumefaciens* mediated genetic transformation in *O. sanctum*. **(A)** binary vector pBI121 with restriction sites and *gusA* with intron; and effect of **(B)** Kanamycin, **(C)** Infection time, **(D)** Acetosyringone, and **(E)** Tween-20 on shoot regeneration frequency in transgenic *O. sanctum*.

### Optimization of parameters affecting transformation efficiency

Genetic transformation in *O. sanctum* was done, using LBA4404 strain of *A. tumefaciens* harboring pBI121 as a binary vector (Fig. 3A). In order to achieve the highest transformation efficiency, several parameters influencing the transformation frequency were optimized. It includes optimization and standardization of effective dose of kanamycin (as above stated), infection time, the concentration of AS, and use of tween-20.

The agrobacterium concentration was previously shown to affect transformation frequencies in many genetic transformation studies. During the present study, *Agrobacterium* suspension of OD600 (0.6) concentration was used to infect the explants. The optimum infection time for achieving highest transient transformation frequency (68.3% ± 0.76) was observed to be 40 min with petiole explants (Fig. 3C). It was found that petioles dipped/treated for 10-30 min in bacterial solution was not optimum as the transient gus expression was very low. However, on further increase in infection time, the transformation frequency was declined drastically (up to 20 %) along with the browning and necrosis of explants. Additionally, it has also been observed that the concentration of AS supplied during transformation study is also a crucial parameter. The established transient expression of gus gene was observed to varied along with acetosyringone (AS) concentration used in co-cultivation period that stimulates the expression of Vir genes in Ti-plasmid. ^34,35^ It was also found that the expression frequency kept on increasing upto 150 μM concentration of AS which gives highest transformation efficiency in *O. sanctum i*.*e*. 85% ± 1.32 (Fig. 3D). With further increase in the concentration of AS the transformation event decreases. The varied concentration from 50-200 μM of AS has been used in plant system depending upon the type plant species. The beneficial effect of AS has been used to maximize the transformation frequencies by many workers.^36,37^ Nonetheless, above stated parameters played an important role in establishing an efficient system for ATMT but the use of Tween-20 during transformation study has also been observed to be highly significant. Surfactant has been used in monocots and dicots to enhance transformation efficiency.^7,38^ By reducing the surface tension, tween-20 encourages bacterium penetration inside the plant cell.^39^ Along with the 150 µM AS and 40 min of infection time, 0.1 % (v/v) of tween-20 was found optimum to achieve highest gus expression in *Ocimum sanctum* (Fig. 3E).

### Regeneration, Gus expression and Molecular analysis

The optimized factors for *in vitro* regeneration [MS medium with BAP (3 mg/L) and NAA (1 mg/L)] from petioles and genetic transformation [(infection medium O.D (600) 0.6 fortified with 150 µM AS + 0.1% (v/v) Tween-20, followed by 40 min of infection and then co-cultivation for 48h in the dark)] worked efficiently in *O. sanctum*. After regeneration, the *in vitro* grown healthy shoots were transferred on the screening medium supplemented with 50 mg/L kanamycin for screening purposes. The regenerated shoots which survived on screening medium was further (1-1.5 cm) transferred to half-strength MS medium for root induction. These putatively transformed plantlets with proper root systems were shifted to glasshouse for further growth.

Meanwhile, after 4-5 days of inoculation on SMM with 50 mg/L kanamycin, healthy and green tissues were assessed for transient gus expression. Gus expression analysis was carried out by histochemical staining with different parts of the *in vitro* regenerated putative transgenic. Distinct blue-stained cells (spots) were obtained indicated the gus expression in various parts of the plants. The expression was maximum at the midrib, veins of the leaves along with petioles, stem, etc. In *O. sanctum*, the GUS-expressing cells displayed as blue-stained small spots, rather than the large blue pattern, similar to experiments done earlier. ^40^ The blue-colored cells were first appeared near veins, but eventually evenly distributed on all sides of the leaf explant, rather than preferentially at the basal end. However, stable gus expression was checked after 3 months (10-12 weeks) of inoculation of putative transgenics surviving on 50 mg/L kanamycin screening medium. The results showed 70% ± 0.35 of stable transformation frequency in *O. sanctum*. The presence of expression was analyzed from randomly selected putatively transformed healthy tissues growing as kanamycin resistant shoots. It was observed that the blue color expression was prominent in leaves, shoot tips, stem, inflorescence, and petioles (Fig. 4A,C,D,E,G,I,J,L). No gus expression was seen in control tissues (Fig. 4B,F,H,K,M).

**Fig. 4.**
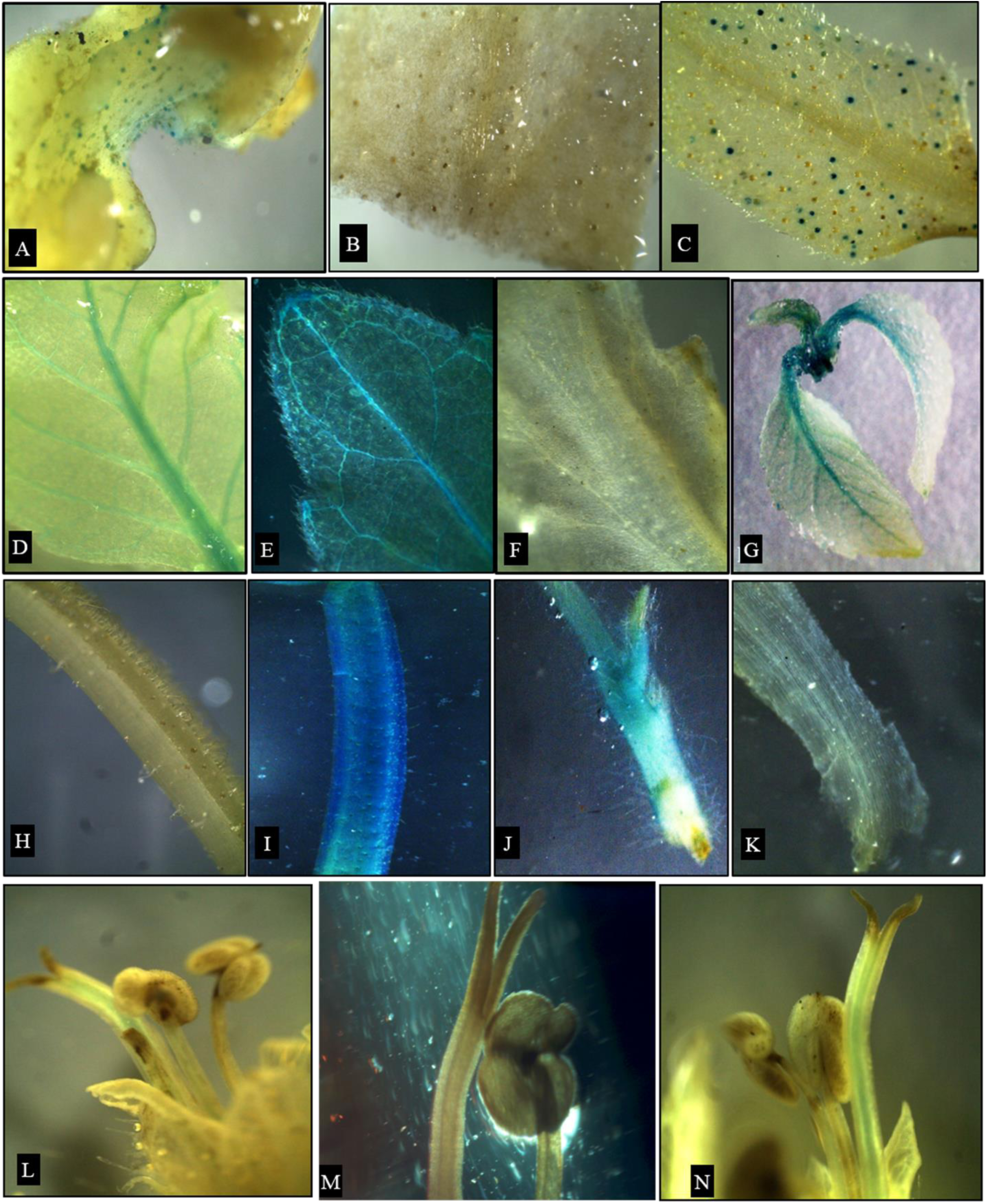
Histochemical Gus expression in *O. sanctum* transgenic plants. (**A**,**C)** transient gus expression in leaf; (**D**,**E**,**G)** gus expression shown in whole transformed leaf and apical node; (**I**,**J)** gus expression in transformed stem and petiole; (**L**,**N)** transformed inflorescence and (**B**,**F**,**H**,**K**,**M)** non-transformed control explants.

In the present study, for the first time, we are describing transient as well as stable gus expression system in *O. sanctum* (*O. tenuiflorum*). In contrast to the previous studies regarding established genetic protocols in *Ocimum basilicum* and *O. citrodonum*; where only transient expression was studied while comparing two different *Agrobacterium* strains GV3101 and EHA105.^8^ Similarly, in *Ocimum gratissimum*, only 20% ± 0.7 of stable transformation frequency was achieved.^7^ Furthermore, in present study, established protocol showed 70% ± 0.35 gus expression frequency. Therefore, the technique is highly efficient and reliable in addition to the fact that there is no ATMT protocol has been reported till date in *O. sanctum*.

In gene transfer technology, transient gus expression is highly advantageous because of the factors like speedy in nature, enabling researchers to get results weekly. However, the stable gus expression is highly recommended for its versatility, simplicity, and robustness. ^41^ Previously many researchers have used transient and stable gus expression as a marker system for assessing the functionality, localization, and transferring of the novel genes. ^42,43^

The genomic DNA from putatively transformed and non-transformed plants was extracted and subjected to PCR analysis. All putative transgenics showed positive results with a band corresponding to approximately 1200 bp and 240 bp for *gus-A* and *npt-II* transgenes, respectively (Fig. 5A,B). Plasmid PBI121 was used as the positive control, and genomic DNA from the wild-type plant was used as the negative control. The results showed the successful integration of *npt-II* and *gus-A* genes in *O. sanctum* transgenic plants.

**Fig. 5.**
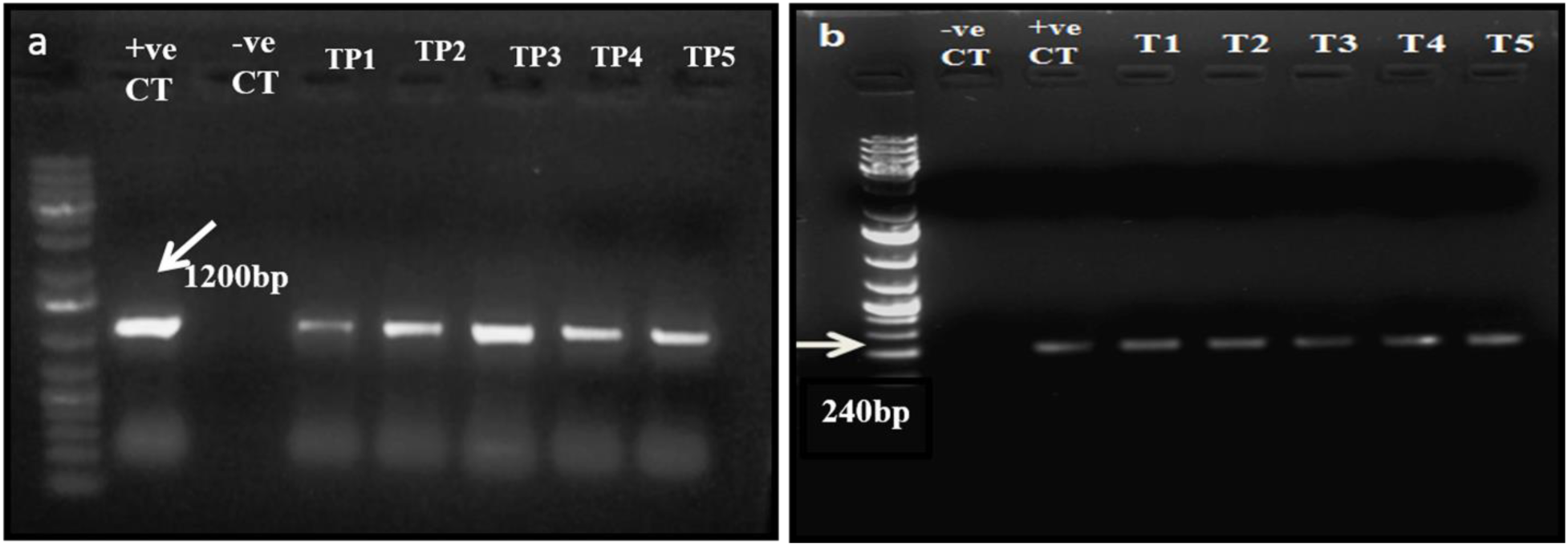
Molecular analysis of transgenic *O. sanctum* plants (**a**) PCR amplification of *gus-A* gene (1200 bp), showing; positive control, negative control and TP1-TP5: gus positive transformants (**b**) PCR amplification of *npt-II* gene (240 bp), positive control, negative control and T1-T5: transformants.

In summary, *Ocimum sanctum* (*O. tenuiliflorum or ‘cim-ayu’*) is a highly rich polyphenolic variety which consists of several bioactive phytoconstituents and therefore, it is very difficult to establish the ATMT protocol. The various factors namely infection time, acetosyringone concentration, use of surfactant etc. showed a great impact on transformation frequency. The optimized factors for *in vitro* regeneration [(MS medium with BAP (3 mg/L) and NAA (1 mg/L)] and genetic transformation [(infection medium O.D.(600) 0.6 fortified with 150 µM AS + 0.1% (v/v) Tween-20, followed by 40 min of infection and then co-cultivation for 48h in the dark)] worked efficiently in genetic transformation of *O. sanctum*. The maximum transformation frequency in *O. sanctum* was 70% ± 0.35. The established method is highly efficient and reproducible *in vitro* protocol for direct regeneration and multiple shoot induction using petiole explant. For the first time, *Agrobacterium tumefaciens* mediated genetic transformation (ATMT) in *O. sanctum* is developed. The protocol can be used for a wide range of genetic and proteomic manipulations and successful integration of desired gene. The technology/protocol provides a powerful tool to modulate the metabolic flux as well as pathway engineering in order to add high pharmaceutical value *in planta*.

## Acknowledgements

Authors wish to express their sincere gratitude to the Director, CSIR-CIMAP, for providing the facilities to carry out this research. The work is supported by grant from the CSIR-CIMAP Network Project BSC-203 (CHEMBIO). SK acknowledge ‘Council of Scientific and Industrial Research (CSIR, New Delhi, India)’ for financial support through CSIR-SRF. The CSIR-CIMAP manuscript number is 2020/JUN/51.

